# Designing a novel Scaffold-Based Multi-Epitope Vaccine to Combat Melioidosis Caused by Burkholderia pseudomallei: An *In-silico* and Immunoinformatics approach

**DOI:** 10.1101/2024.06.18.599592

**Authors:** Shakilur Rahman, Ardhendu Das, Amit Kumar Das, Ditipriya Hazra, Amlan Roychowdhury

## Abstract

*Burkholderia pseudomallei*, the gram-negative bacteria causing melioidosis, is becoming a serious threat to healthcare settings. In recent years, *B. pseudomallei* has been identified as an emerging and significant etiological agent responsible for localized pyogenic infections primarily observed in India and South Asia. At present, no vaccine against melioidosis is available in the treatment system. This study has undertaken an *in-silico* reverse vaccinology approach to design a novel multi-epitope vaccine for treating *B. pseudomallei*-mediated infections. B-cell and T-cell epitopes have been predicted and stitched to develop a multi-epitope vaccine. The predicted vaccine is found to be non-toxic, non-allergic, and immunogenic in nature. Immune simulation results indicate that the designed vaccine can generate an immune response resembling a real-life scenario. The 610 amino-acid long vaccine construct has been codon-optimized and could be cloned in the E. coli K12 system. These findings from this immunoinformatics study offer a foundation for developing a tailored, safe, and potent vaccine targeting *B. pseudomallei*.

## 1. INTRODUCTION

*Burkholderia pseudomallei* is an aerobic, Gram-negative soil-dwelling bacillus and the causative agent of melioidosis [1]. Melioidosis, also recognized as Whitmore’s disease, presents a significant concern as it is capable of infecting both animal and human populations [2]. *B. pseudomallei* can survive in the soil for many years & when contaminated particles or aerosols are inhaled, swallowed, or exposed to wounds, melioidosis occurs [3]. Typically, it leads to the formation of abscesses in the lung, liver, spleen, skeletal muscle and parotid glands and becomes severe among immunocompromised patients, particularly people who have previous health issues like diabetes, chronic renal failure, thalassemia, etc.[4]. According to a study on spatial modeling conducted in 2016, there were around 165,000 instances of melioidosis globally in humans, of which 89,000 (or 54%) resulted in mortality [5].

*B. pseudomallei* utilizes several virulence factors like quorum sensing, secretion system, capsular polysaccharide, flagella, etc., along with siderophores and different enzymes like protease, lipase, catalase, superoxide dismutase, etc., for disease colonization, persistence, and latency [6,7]. Being a facultative intracellular pathogen, the life cycle of *B. pseudomallei* includes adhesion, penetration into host cells, phagosome escape, cytosolic replication, and cell-to-cell dissemination [8]. Adhesion is an essential step in the establishment of an infection, allowing the bacteria to interact and attach to host tissues [9]. Cell adhesion in *B. pseudomallei* is mediated by certain membrane proteins, like flagellin (fliC) and extracellular adherence proteins[10]. Trimeric autotransporter adhesin (BpaC) is essential for the pathogen’s initial attachment and mediating adherence to respiratory epithelial cell types [11]. After attachment, *B. pseudomallei* often uses the secretion system type III to infiltrate epithelial cells. It also helps to replicate in the cytoplasm by escaping from the phagosome [12,13]. Similar to BimA-mediated actin polymerization of the host cell, hemolysin activator-like protein precursor (fhaC) functions as an effector and allows *B. pseudomallei* to migrate freely once it has entered the cytoplasm [14,15]. Certain outer membrane receptors that engage with the inner membrane TonB complex are necessary for *B. pseudomallei* to absorb ferric siderophore complexes. It is used to accumulate iron from the host to proliferate in iron-deficient environments [16,17]. The oligopeptide-binding protein A (OppA), a part of the ATP binding cassette (ABC) transport system, is known for its key role in nutrient uptake and recycling of cell-wall peptides. OppA helps in bacterial survival, virulence and overall pathogenicity [18]. Upon initial pathogen exposure, the innate immune system identifies conserved surface motifs known as pathogen-associated molecular patterns (PAMPs), for example, lipopolysaccharide (LPS), type III secretion system, flagellin, peptidoglycan, etc., through host-cell pattern-recognition receptors, particularly Toll-like receptors (TLRs) [19,20]. These TLRs then trigger the innate immune response, establishing a vital connection between innate and adaptive immunity. Besides, TssM is a secreted deubiquitinase that inhibits TLR-mediated NF-κB activation by ubiquitinating signaling intermediates, therefore suppressing the innate immune responses [21]. TLR2 is activated in the host by exposure to *B. pseudomallei* lipopeptides and peptidoglycan, which is followed by TLR1 or TLR6 [22]. Multiple studies support the induction of reactive T-cells in humans against *B. Pseudomallei* infection, and the findings show that the CD4^+^ and CD8^+^ T cells are activated and able to produce specific IFNγ in response to antigens, demonstrating the immune system’s readiness to fight infections [23–25]. Host-pathogen interactions take place in the bacterial membrane, and the antigenic proteins harbouring the extracellular, periplasmic, and outer membrane are exposed to the host’s immune surveillance; these proteins emerge as potential vaccine candidates in the battle against infectious diseases like Melioidosis [26,27].

Conventional vaccinology uses time-consuming and expensive approaches that include growing bacteria for pathogenic detection, isolation, characterization, reinjection, and inactivation in the host to elicit an immune response [28]. “Reverse Vaccinology” eliminates the need for culture-specific microorganisms by systematically developing vaccines employing genetic data and computational tools [29,30]. In a multi-epitope vaccine, multiple epitopes from different antigenic proteins are used and it results in a complete immune response against the pathogen [31]. Utilizing this method, both humoral and cell-mediated immune responses can be achieved by the incorporation of immunodominant conserved epitopes [32]. Reverse vaccinology approach has been proposed to design vaccines against different pathogenic bacteria, like *Staphylococcus aureus* [33], *Listeria monocytogenes* [34], *Klebsiella aerogenes* [35], *Porphyromonas gingivalis* [36].

In this current study, subtractive proteomics and experimental data coupled with scaffold-based protein engineering, molecular dynamics simulation and immunoinformatic approaches have been used to develop a multi-epitope vaccine (MEV) against *B. pseudomallei.* The scaffold-based construction of engineered MEV has been applied for the very first time against *B. pseudomallei*.

## 2. RESULTS

### 2.1. Retrieval of sequence and antigenicity of proteins

An extensive literature survey identified several proteins implicated in the virulence or survival of *B. pseudomallei*. During the selection of potential vaccine candidates, preference was given to proteins deemed essential or associated with virulence, with further criteria including the antigenicity score above 0.4 (Table 1). The amino acid sequences of the proteins Periplasmic oligopeptide-binding protein (OppA), Flagellin (fliC), Porin related exported protein, Trimeric autotransporter adhesin (BpaC), Porin protein (OpcP), Hemolysin-related protein, Iron complex outer membrane receptor protein (TonB), Type III secretion system effector protein (BapA), hemolysin activator-like protein precursor (FhaC), Membrane protein insertase (YidC), Type III secretion system effector protein (BapC), Regulator and type III secretion system effector protein (BbrD), deubiquitinase (TssM), OstA-like protein (LptA) were obtained from UniProt database and the Burkholderia Genome Database. The selected proteins were found to be antigenic as the antigenicity values provided by VaxiJen v2.0 met the threshold value.

**Table 1.** List of proteins considered for vaccine designing, along with their antigenicity scores.

| *Protein* | *Name* | *Localization* | *UniProt ID* | *Antigenicity Score* |
| --- | --- | --- | --- | --- |
| BPSS2141 | Periplasmic oligopeptide-binding protein, **OppA** | Periplasmic | Q63ID0 | 0.48 |
| BPSL3319 | Flagellin, **FliC** | Extracellular | H7C7G3 | 0.64 |
| BPSS1679 | Porin related exported protein | Outer Membrane | Q63JN8 | 0.62 |
| BPSL1631 | Trimeric autotransporter adhesin, **BpaC** | Outer Membrane | Q63UH1 | 1.28 |
| BPSS0879 | Porin protein, **OpcP** | Outer Membrane | Q63LY3 | 0.66 |
| BPSL1661 | Putative hemolysin-related protein, | Hemolysin-related protein | Q63UE4 | 0.77 |
| BPSS1850 | Iron complex outer membrane receptor protein, **TonB** | Outer Membrane | Q63J68 | 0.68 |
| BPSS1728 | Hemolysin activator-like protein precursor, **FhaC** | Outer Membrane | Q63JI8 | 0.52 |
| BPSS1528 | Type III secretion system effector protein, **BapA** | Extracellular | Q63K38 | 0.77 |
| BPSL0078 | Membrane protein insertase **YidC** | Cytoplasmic Membrane | Q63YW1 | 0.66 |
| BPSS1526 | Type III secretion system effector protein, **BapC** | Periplasmic / Extracellular | Q63K40 | 1.6 |
| BPSS1521 | Regulator and type III secretion system effector protein, **BbrD** | Cytoplasmic / Extracellular | Q63K45 | 1.34 |
| BPSS1512 | Deubiquitinase, **TssM** | Extracellular | Q63K53 | 1.28 |
| BPSL0535 | OstA-like protein (lipopolysaccharide export system protein), **LptA** | Extracellular | Q63XK5 | 1.31 |

### 2.2. Predicted Linear B-cell epitopes

Using ABCpred, B-cell epitopes from the aforementioned proteins were determined; peptides with a score of more than 0.85, which denotes strong immunogenicity, were chosen for further examination. B-cell epitopes with the sequence, antigenicity scores, allergenicity and toxicity predictions are shown in Table 2. From the set of predicted epitopes, the following two epitopes, YQHASGTQRVDATTTQ (BPSS1679) and TATGTDSTASGSNSTA (BPSL1631), were used in multi-epitope vaccine (MEV) construction. Along with these two top-scoring predicted epitopes, three experimentally obtained B-cell epitopes from BPSL3319 were also used in the MEV construction (Table 5).

**Table 2.**
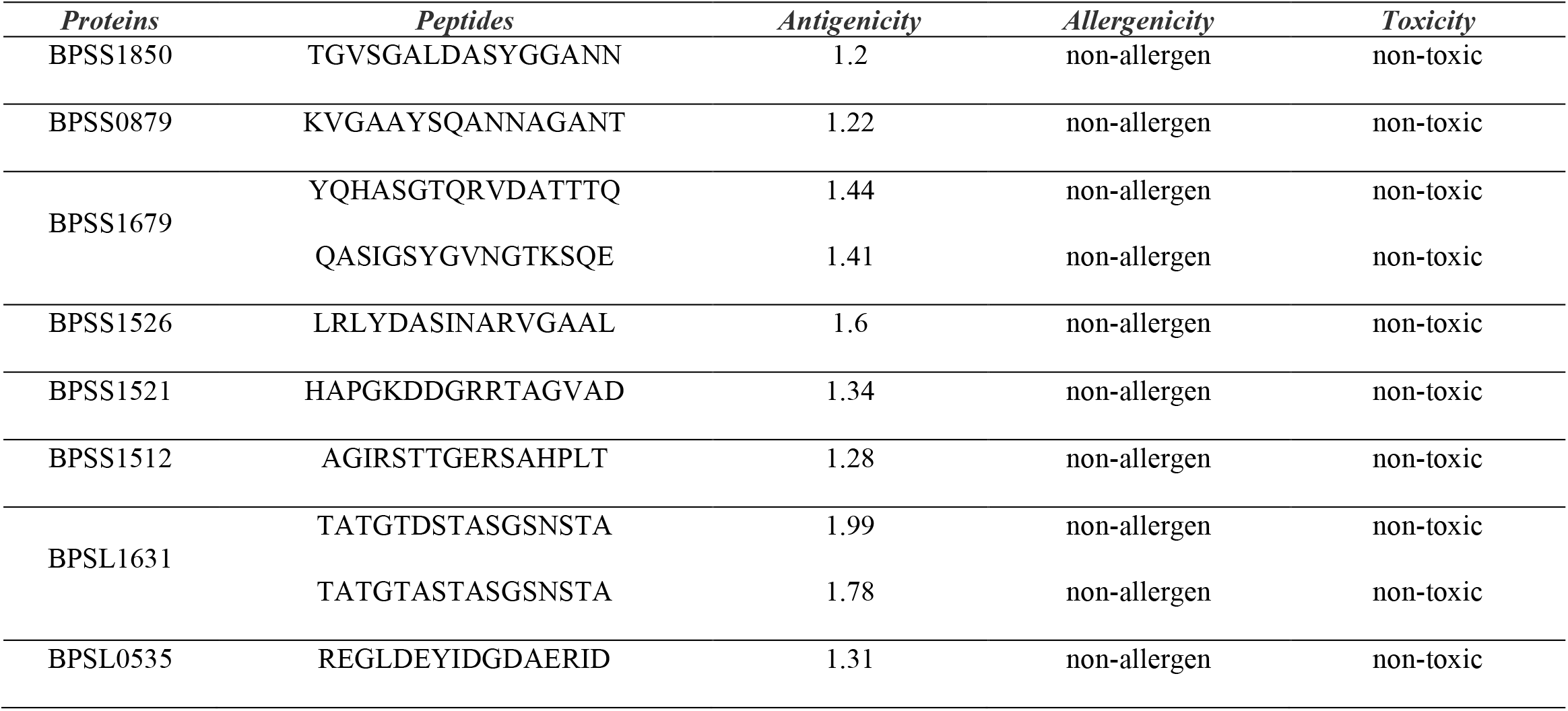
List of the predicted B-cell epitopes.

### 2.3. Predicted T-cell epitopes

The antigenic proteins mentioned above were subjected to CTL (cytotoxic T lymphocyte) and HTL (helper T lymphocyte) epitope prediction, and the results are depicted in Tables 3 and 4. From the predicted epitopes, 6 CTL epitopes and 6 HTL epitopes, along with an experimentally obtained potential T-cell epitope (KGGFTFTDQNNQALSST) from **BPSL3319** protein were selected for the final construction of MEV (Table 5).

**Table 3.**
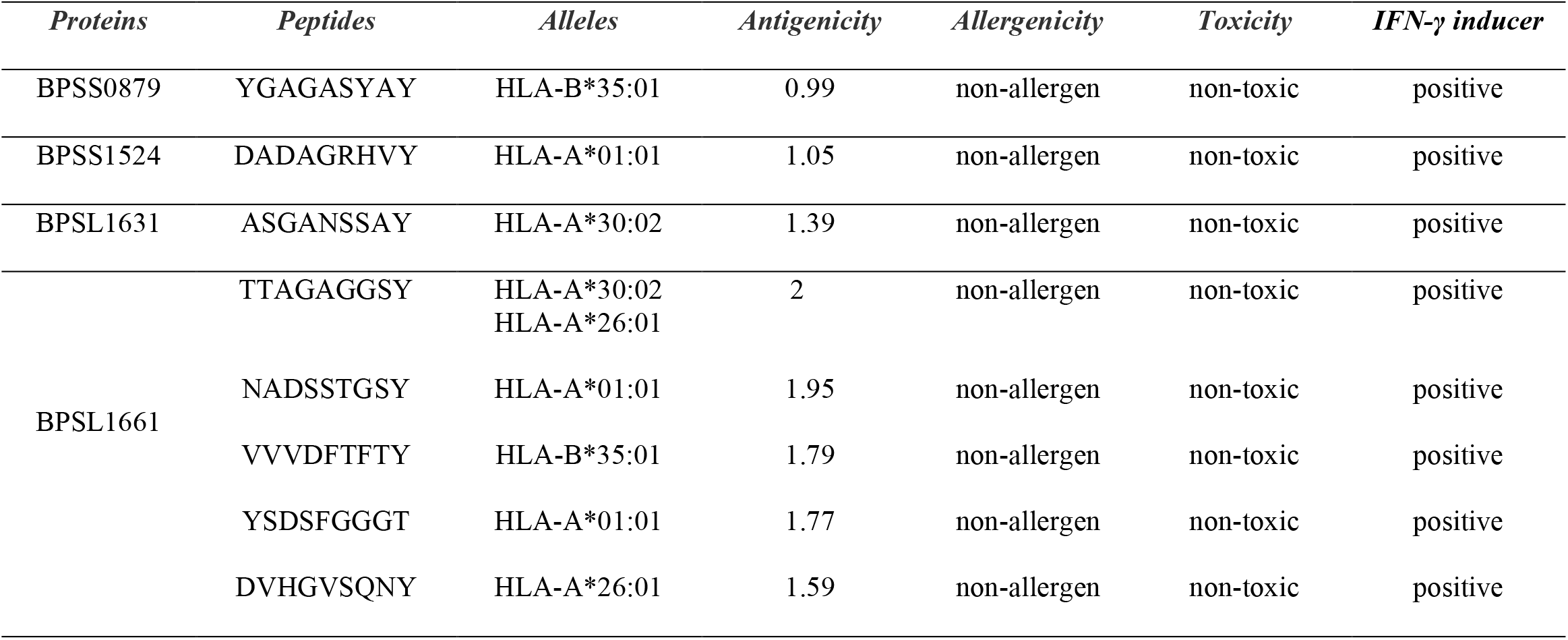
List of predicted CTL epitopes.

**Table 4.** List of predicted HTL epitopes.

| *Proteins* | *peptides* | *Alleles* | *Antigenicity* | *Allergenicity* | *Toxicity* | *IFN-γ inducer* |
| --- | --- | --- | --- | --- | --- | --- |
| BPSS1850 | SAGARIEHVRLDPSA | HLA-DRB4*01:01 | 1.58 | non-allergen | non-toxic | positive |
| BPSS0879 | EANVKYNLTPALGLG | HLA-DRB1*07:01 | 1.23 | non-allergen | non-toxic | positive |
| BPSS1728 | YLRINRNPNAANANG | HLA-DRB3*02:02 | 1.19 | non-allergen | non-toxic | positive |
| BPSS1528 | AGGLERDAPGERAAG | HLA-DRB1*03:01 | 1.34 | non-allergen | non-toxic | positive |
| BPSL1661 | SDGLTKDSRVTISGT | HLA-DRB1*03:01 | 1.15 | non-allergen | non-toxic | positive |
|  | PSDGLTKDSRVTISG | HLA-DRB1*03:01 | 1.15 | non-allergen | non-toxic | positive |
| BPSL0078 | YVIGVDTKIENVGAA | HLA-DRB1*03:01 | 1.31 | non-allergen | non-toxic | positive |
|  | RGSYVIGVDTKIENV | HLA-DRB1*03:01 | 1.23 | non-allergen | non-toxic | positive |

**Table 5.**
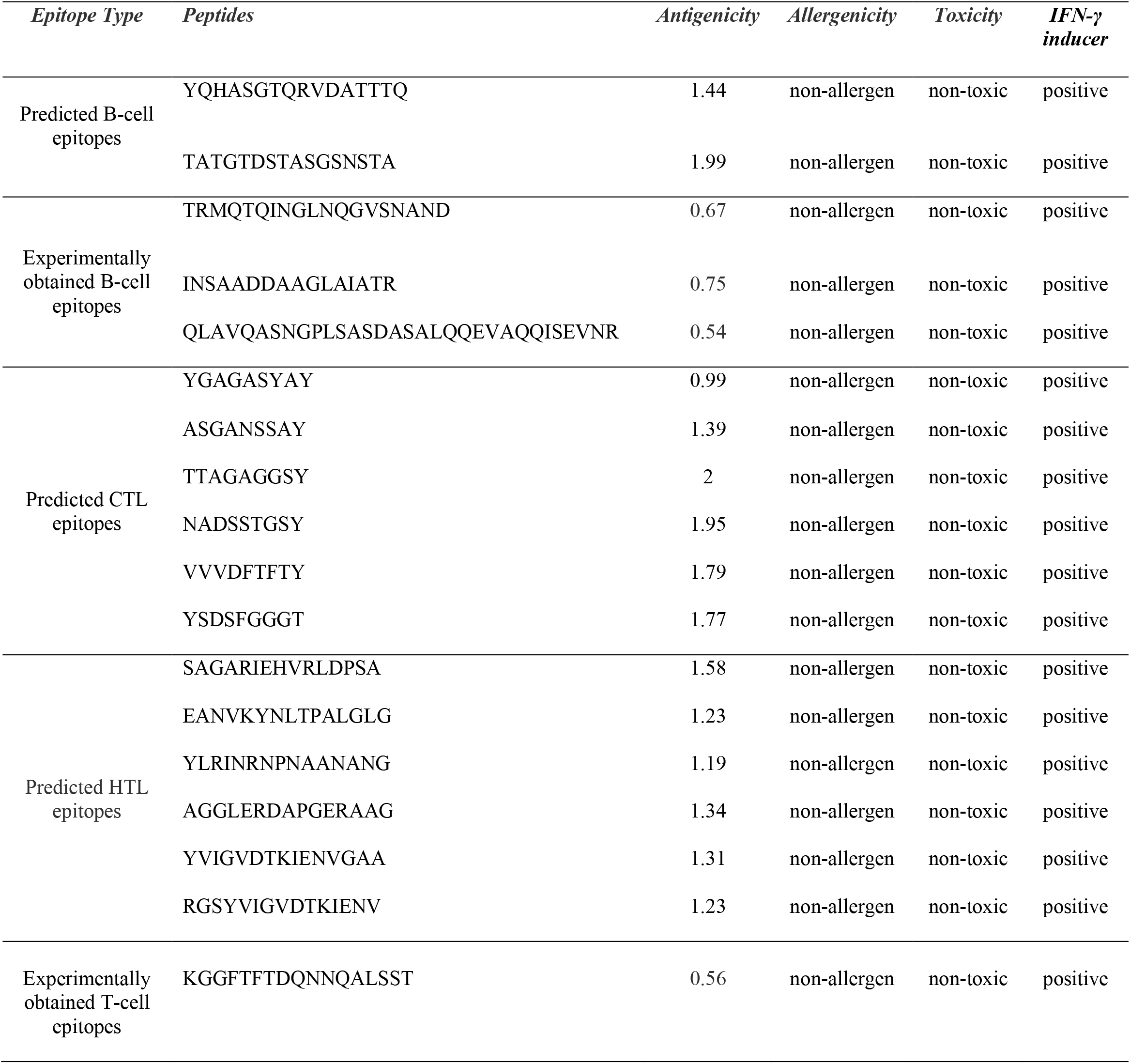
List of epitopes selected for MEV construction.

| *Epitope Type* | *Peptides* | *Antigenicity* | *Allergenicity* | *Toxicity* | *IFN-γ inducer* |
| --- | --- | --- | --- | --- | --- |
| Predicted B-cell epitopes | YQHASGTQRVDATTTQ | 1.44 | non-allergen | non-toxic | positive |
|  | TATGTDSTASGSNSTA | 1.99 | non-allergen | non-toxic | positive |
| Experimentally obtained B-cell epitopes | TRMQTQINGLNQGVSNAND | 0.67 | non-allergen | non-toxic | positive |
|  | INSAADDAAGLAIATR | 0.75 | non-allergen | non-toxic | positive |
|  | QLAVQASNGPLSASDASALQQEVAQQISEVNR | 0.54 | non-allergen | non-toxic | positive |
| Predicted CTL epitopes | YGAGASYAY | 0.99 | non-allergen | non-toxic | positive |
|  | ASGANSSAY | 1.39 | non-allergen | non-toxic | positive |
|  | TTAGAGGSY | 2 | non-allergen | non-toxic | positive |
|  | NADSSTGSY | 1.95 | non-allergen | non-toxic | positive |
|  | VVVDFTFTY | 1.79 | non-allergen | non-toxic | positive |
|  | YSDSFGGGT | 1.77 | non-allergen | non-toxic | positive |
| Predicted HTL epitopes | SAGARIEHVRLDPSA | 1.58 | non-allergen | non-toxic | positive |
|  | EANVKYNLTPALGLG | 1.23 | non-allergen | non-toxic | positive |
|  | YLRINRNPNAANANG | 1.19 | non-allergen | non-toxic | positive |
|  | AGGLERDAPGERAAG | 1.34 | non-allergen | non-toxic | positive |
|  | YVIGVDTKIENVGAA | 1.31 | non-allergen | non-toxic | positive |
|  | RGSYVIGVDTKIENV | 1.23 | non-allergen | non-toxic | positive |
| Experimentally obtained T-cell epitopes | KGGFTFTDQNNQALSST | 0.56 | non-allergen | non-toxic | positive |

### 2.4. Multi Epitope Vaccine (MEV) Construction, Modelling and validation

The MEV was constructed using 5 B-cell epitopes and 13 T-cell epitopes, summarized in Table 5. The truncated OppA from *B. pseudomallei* was used as the scaffold. Predicted B-cell epitopes were joined using linker KK, CTL epitopes were joined using AAY linker, and HTL epitopes were joined using GPGPG linker. Human β-defensin-3 was used as an adjuvant and linked to the N terminus end of the protein to form the complete MEV comprised of 610 amino acids. The 3D structure of the constructed MEV (Figure 2) was predicted using Robetta. After validating with the PROCHECK, 90.2% of the amino acids of the predicted model were found in the most favoured region, 7.5% in the additionally allowed and 1.3% in the generously allowed region (Supplementary). The flexibility of the designed MEV structure was indicated by the top 10 superimposed models (Figure 3). The maximum RMSF value was 4.54 Å, and the minimum RMSF value was 0.075 Å. The peaks indicated the area with the highest flexibility. Moreover, both terminals show flexibility under accepted ranges.

**Figure 1.**
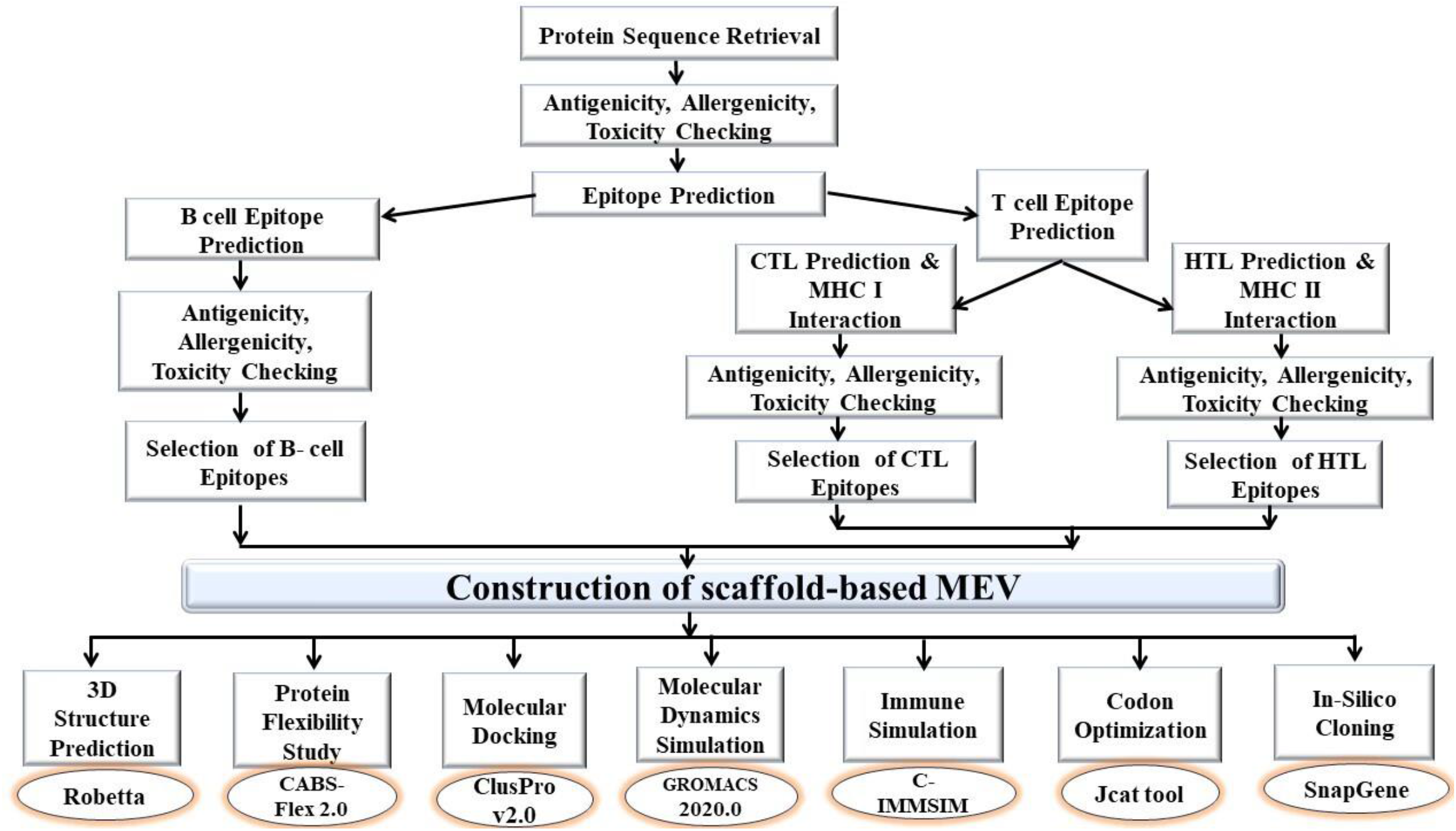
Flowchart representation of the overall workflow to design the *in-silico* scaffold-based Multi-Epitope Vaccine (MEV) against *Burkholderia pseudomallei*.

**Figure 2.**
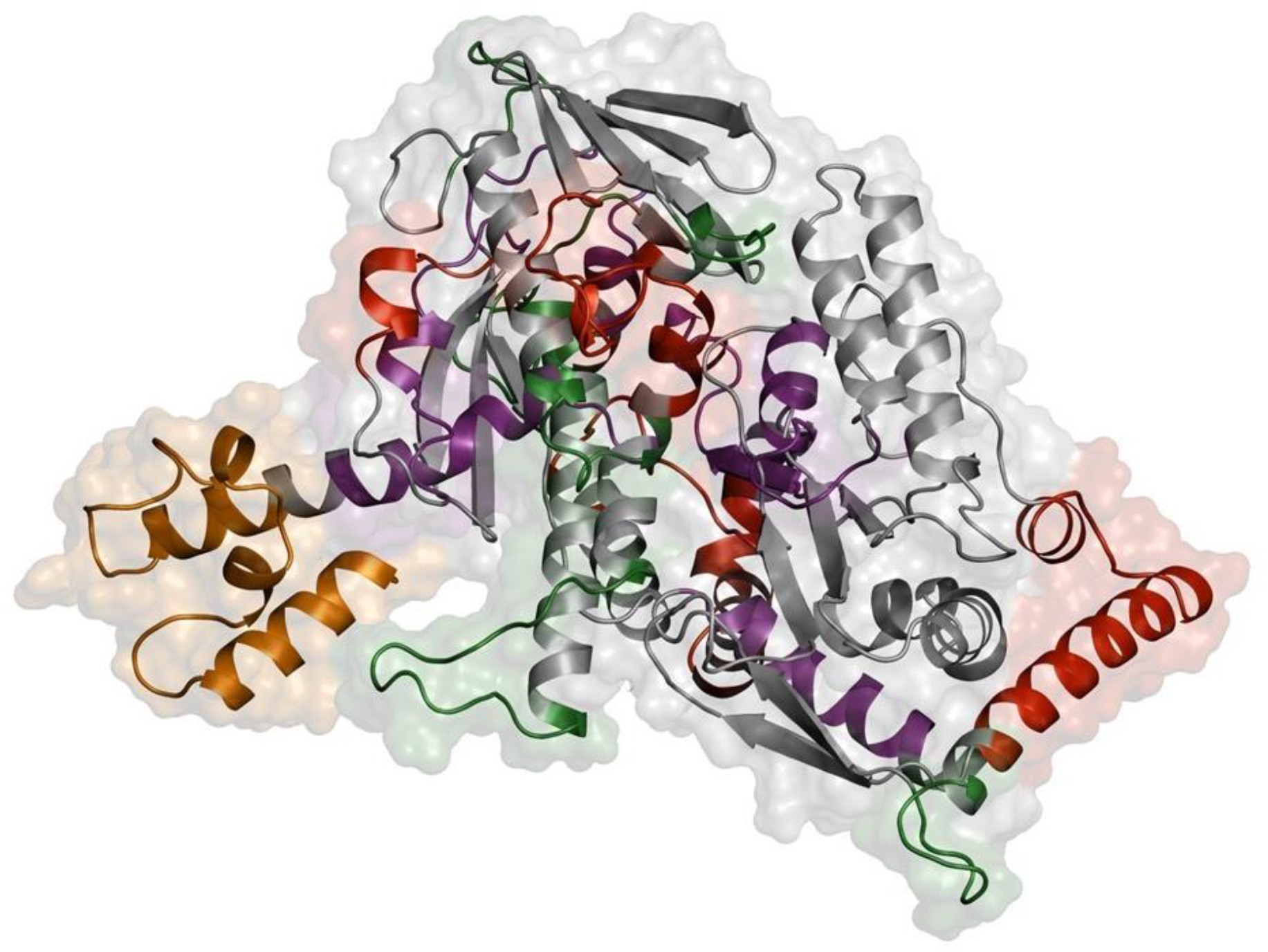
The tertiary structure of the MEV is represented in the cartoon version with an essence of the surface. The B-cell, cytotoxic T-cell and Helper T-cell epitopes are depicted in red, green, and purple colours, respectively. The adjuvant is represented in orange colour and the scaffold protein is in grey colour.

**Figure 3.**
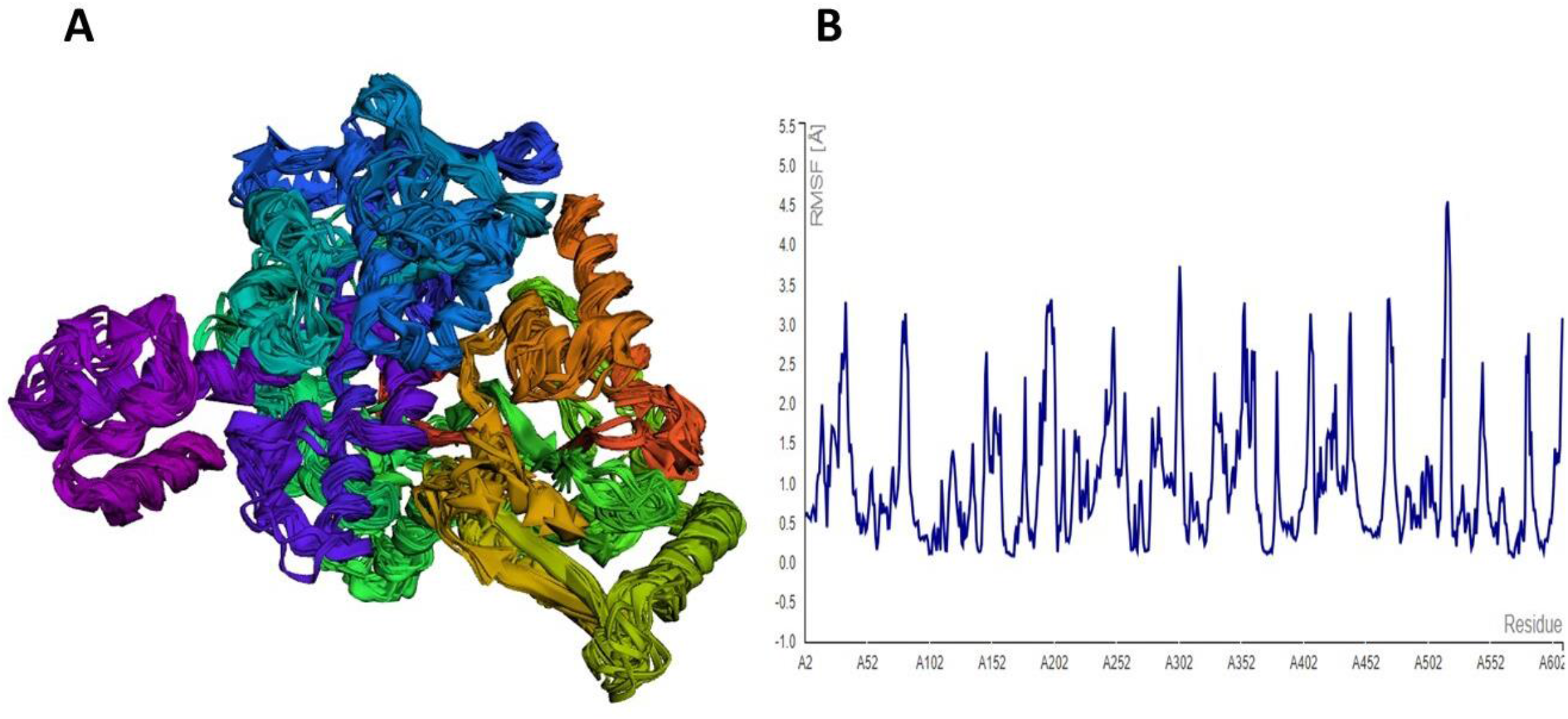
Analysis of structural stability of the MEV. (A) Representation of top 10 superimposed models and (B) RMSF of the MEV obtained from CABSFLEX 2.0.

### 2.5. Study of MEV-TLR2 interaction

The first pose (cluster_0) for the MEV-TLR2 complex was chosen as the best one on the basis of the docking score. The Molecular Mechanics/Generalized Born Surface Area (MM/GBSA) method using the HawkDock platform has shown a favourable binding energy of −109.23 kcal/mol for the MEV-TLR2 complex. The residues from both MEV and TLR2 that significantly contributed to the formation of the MEV-TLR2 complex are listed in (Figure 6) along with the individual binding free energy expressed in kcal/mol.

### 2.6. Studying conformational dynamics of MEV & MEV-TLR2 complex

The classical molecular dynamics simulation approach was applied to both systems to understand the conformational dynamics and structural stability of the MEV and the MEV-TLR2 complex. The stability of the system was depicted by the Root-Mean-Square Deviation (RMSD); on the other hand, the fluctuations of the atoms and residues of the system were described by the Root-Mean-Square Fluctuation (RMSF). In our study, an average RMSD of 0.51 nm and 0.32 nm was observed for MEV and the MEV-TLR2 complex, respectively (Figures 4 & 5). The average RMSF of 0.19 nm was observed for the MEV (Figures 4 & 5). The average R_g_ values were found to be 2.69 nm and 3.75 nm for the MEV and the MEV-TLR2 complex, respectively, and remained stable throughout the simulation, which proves the compactness and structural integrity of both systems.

**Figure 4.**
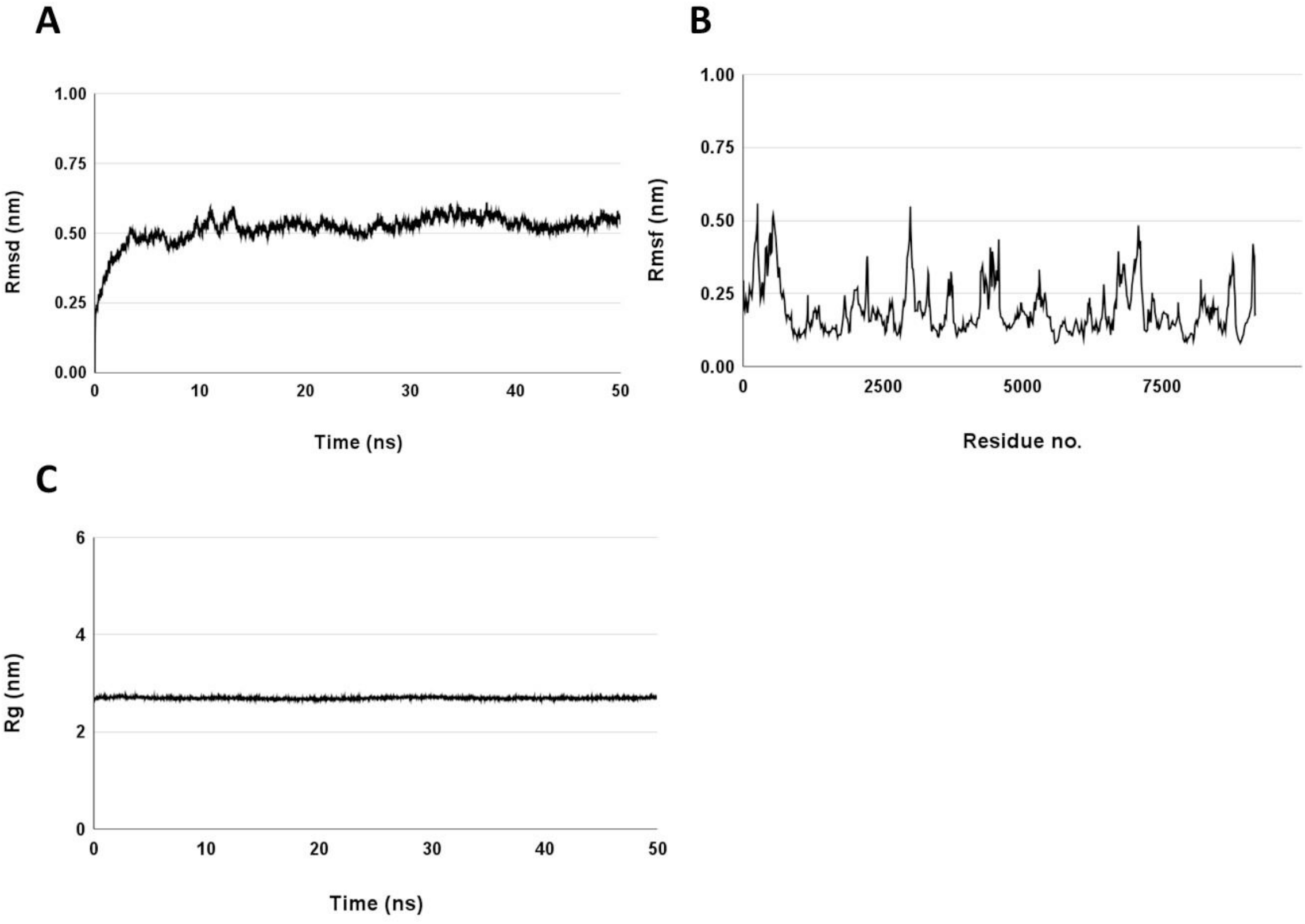
The stability of the MEV structure is indicated by the MD simulation study. (A) RMSD analysis shows a deviation of 0.51 nm, (B) RMSF is observed within an allowed range with some fluctuating residues and (C) Rg of the MEV.

**Figure 5.**
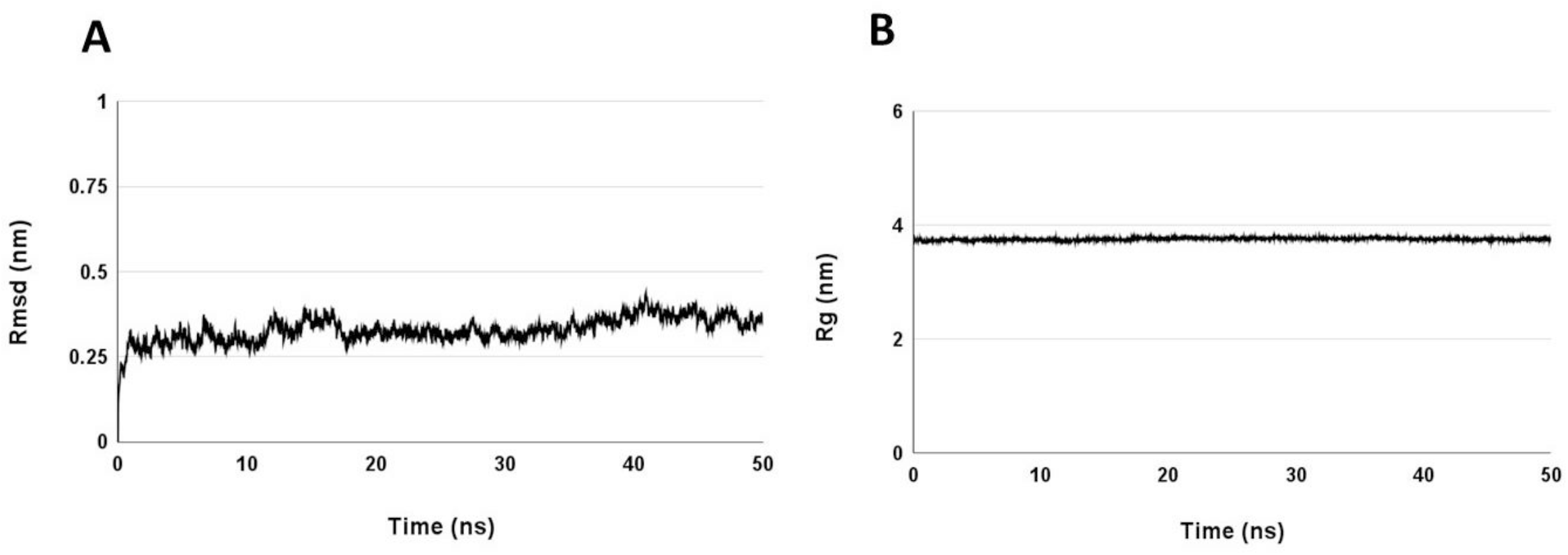
The stability of the MEV-TLR2 complex structure is indicated by the MD simulation study. (A) RMSD analysis shows a deviation of 0.32 nm and (B) Rg of the MEV-TLR2 complex.

**Figure 6.**
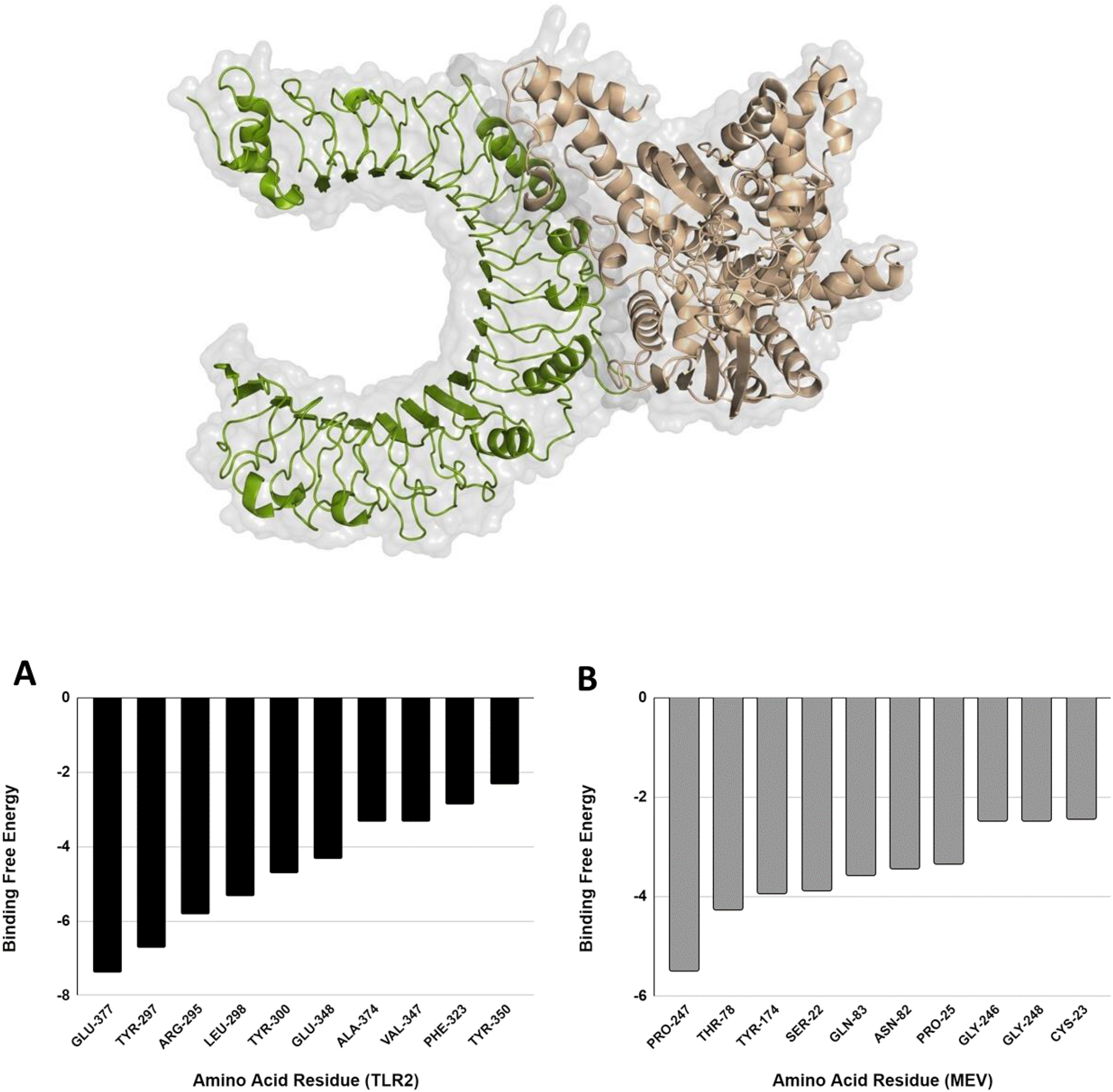
The tertiary structure of the MEV-TLR2 complex is represented in the cartoon version with an essence of the surface. (A) The amino acid residue-wise binding free energy calculation for TLR2 and (B) The amino acid residue-wise binding free energy calculation for MEV.

### 2.7. Analysis of immune simulation of MEV

Immune simulation analysis showed that the population of IgG and IgM antibodies increased gradually after the initial injection. Subsequent to the third and second booster injections, there was a further increase in the population of IgG and IgM antibodies. After every booster shot, the concentration of B-cells increased noticeably, and they were mainly comprised of active B-cells (Figure 7-A, C). The total number of plasma cells increased with every booster dose. Most plasma cells released IgM antibodies after the second booster. But with the third dose, there was a more balanced production, with almost equal amounts of IgM and IgG antibodies released (Figure 7-B). With every booster dose, the number of helper T cells increased by three times. Following the third booster dose, memory T_H_ cell concentration reached its maximum level (Figure 7-E). On the other hand, throughout the simulation, the quantity of natural killer cells and cytotoxic T cells stayed constantly high. IFN-γ production was markedly increased during this phase, and other cytokines, including TGF-β, IL10, and IL12, were also elevated (Figure 7-D).

**Figure 7.**
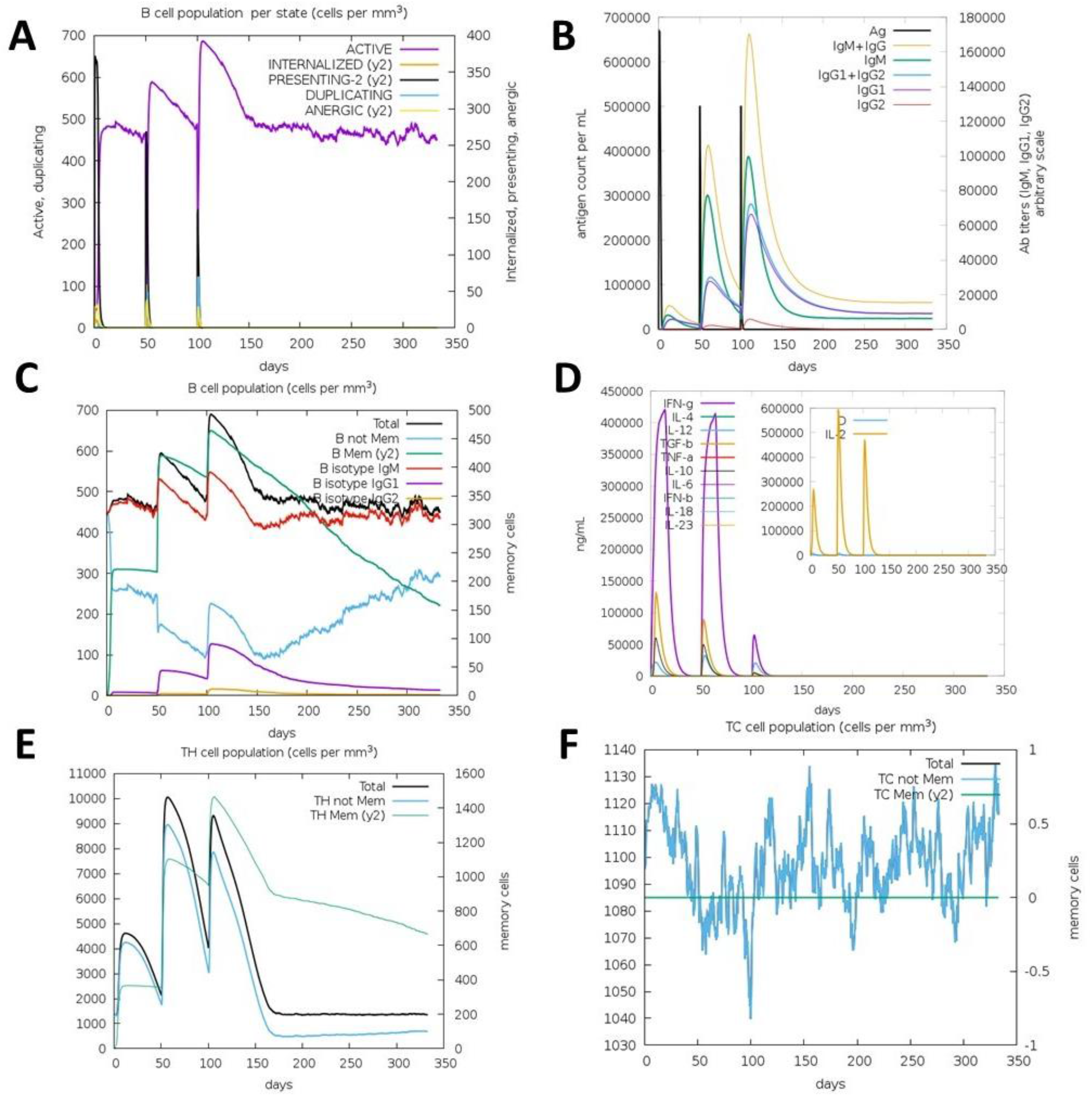
Analysis of *in-silico* Immune Simulation study. (A) The population of B-cells per state, (B) Increased levels of IgM and IgG antibodies produced by B cells upon antigen exposure, (C) B-cell population per mm^3^ in a timeframe of 350 days after cytokines induction, (D) Increased concentration of cytokines like IFN-γ, TGF-β, IL-10 and IL-12 after vaccine administration, (E) Population of Helper T-cells per mm^3^ in a timeframe of 350 days after cytokines induction and (F) Population of cytotoxic T-cells per mm^3^ in a timeframe of 350 days after cytokines induction.

### 2.8. Codon optimization & *in-silico* cloning

The 1830 bp sequence having a 0.991 CAI value with 52.62% GC content was generated after codon optimization of the MEV construct and it suggested that the protein could be expressed in the K12 strain of *E. coli.* Cloning of the optimized DNA and vaccine gene into the ‘Multiple Cloning Sites’ (MCS) of the pET-28a (+) cloning vector was done *in-silico*, keeping the *BamHI* and *XhoI* at the 5’ and 3’-terminals of the DNA sequence, respectively (Figure 8). The predicted solubility score of the MEV was found to be 0.927 in the *E. coli* expression system, and it indicated that the MEV was highly soluble.

**Figure 8.**
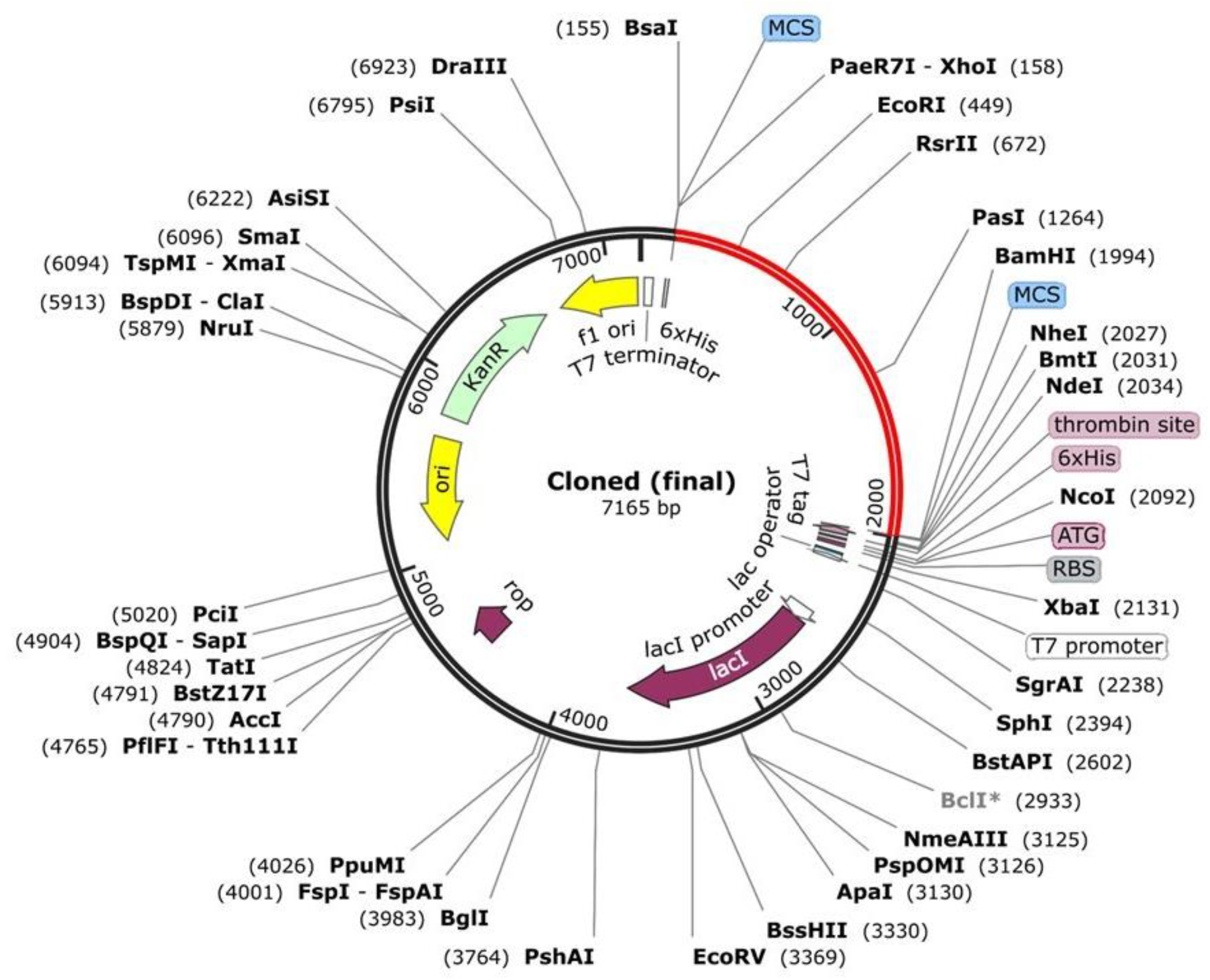
Representation of *in-silico* cloning of the MEV into pET28a (+) vector. The cloned vaccine construct is highlighted in red colour.

## 3. DISCUSSION

There are very few treatment options available since clinical data indicate that all *B. pseudomallei* species are resistant to a variety of antibiotic classes, and some isolates are even resistant to front-line medications like ceftazidime and trimethoprim-sulfamethoxazole (TMP-SMX) [37]. Hence, developing new therapeutics against *B. pseudomallei* infection has become an active area of research [38,39].

In this study, several antigenic proteins were evaluated for the vaccine preparation, as they have fundamental roles in adhesion and penetration into host cells, phagosome escape, cytosolic replication, and cell-to-cell dissemination, which are all critical phases of the bacterial life cycle (Table 1). The epitopes from the highly immunogenic proteins were predicted using an immunoinformatic approach and subtractive epitope technique (Table 5). It is also advantageous to include an adjuvant in the vaccine formulation because it can stimulate cell-mediated immunity, boost a strong immune response, facilitate mucosal transport of vaccines, and lower the frequency of vaccination or antigen dosage [40,41]. In mammalian immunity, adjuvant β-Defensin 3 (45 amino acids long) is known to have antimicrobial and chemoattractant effects [42,43]. For this reason, β-Defensin 3 has been used as an adjuvant during the MEV construction. The B-cell, CTL and HTL epitopes were linked using KK, AAY, and GPGPG linkers, respectively. Further, an EAAAK linker was used to link the β-Defensin 3 adjuvant. The molecular weight of the designed vaccine was 65.60 KDa. As revealed in the previous studies, TLR2 plays a crucial role in host-pathogen interaction in *B. pseudomallei* infections; the molecular interaction between MEV and TLR2 (Figure 6), and the stability of MEV-TLR2 complex was validated by MD simulation.

T-cell stimulation by antigen-presenting cells is necessary for a vaccination to be effective [44]. T-lymphocytes are also essential for the longevity and cross-reactivity of vaccinations [45]. Upon administration of the vaccine to a patient, the activation of the Helper T-cell population has been noticed [46]. With the help of IFN-γ, TNF-α, and other cytokines, the activated Helper T-cells further activate the B-cell population [47]. In response, immunoglobulins are produced by the induced B-cells, more precisely the IgG immunoglobulins [48]. The immune simulation data showed that the active Helper T-cell populations increased significantly after vaccination, along with the rise of cytokines IFN-γ and TNF-α. Further, these cytokines induced the B-cell population, resulting in an increased level of IgG antibodies. The first induction of memory B-cell population and a constant rise in cytokines such as IFN-γ, TGF-β, IL-10, and IL-12 following vaccination injection demonstrate that the developed MEV construct exhibits congruence with real-life immune responses. Multi-epitope vaccines are highly specific, elicit targeted immune responses, and prevent allergic reactions; on the other hand, epitope-based peptide vaccines have particular challenges due to their purity as well as stability, degradation in the serum, and lack of post-translational modifications [49–51]. The results of this comprehensive study demonstrated that our developed vaccine architecture may elicit an immune response without causing an allergic reaction. Consequently, animal models and clinical trials must be conducted to assess the effectiveness of the multi-epitope vaccination in the host.

## 4. CONCLUSION

Melioidosis, prevalent in tropical regions, is a major public health concern due to its high mortality rates, with the potential for epidemic spread beyond endemic areas, raising global health security concerns. This study explores bacterial antigens to provide a scaffold-based multiepitope vaccine against *B. pseudomallei*. The reverse vaccinology approach provides potential B-cell and T-cell epitopes, further stitched into a 65.60 kDa muti-epitope vaccine construct. The predicted MEV is found to be non-toxic, non-allergic, and, most importantly, highly antigenic. The designed MEV maintains a stable interaction with Toll-like receptor 2. Regulation of lymphocytes and cytokines after vaccine administration resembles the real-life immune response scenario. Finally, the codon-optimized 1830 bp-long MEV construct can be cloned and expressed in the *E. coli* K12 bacterial system. Additional experimental trials are essential to confirm the efficacy of the scaffold-based MEV constructs against *B. pseudomallei*.

## 5. MATERIALS & METHODS

### 5.1. Selection of Antigen & Retrieval of Sequence

The protein sequences were obtained from the UniProt database (https://www.uniprot.org/) and the Burkholderia Genome Database (https://www.burkholderia.com/). The selected proteins were subjected to antigenicity prediction using VaxiJen v2.0 (http://www.ddg-pharmfac.net/vaxijen/VaxiJen/VaxiJen.html) [52], taking 0.4 as the threshold value.

### 5.2. Prediction of B-Lymphocyte epitopes

B-cell epitopes are crucial for generating host antibody response. The linear B-cell epitopes were predicted using an Artificial neural network-based B-cell epitope prediction server - ABCpred server (https://webs.iiitd.edu.in/raghava/abcpred/index.html) [53] with a cut-off score of 0.51. Then, the predicted B-cell epitopes were further subjected to antigenicity checking using VaxiJen v2.0 (http://www.ddg-pharmfac.net/vaxijen/VaxiJen/VaxiJen.html), taking 0.4 as the threshold value. For the allergenicity screening, the AllerTop v2.0 server (https://www.ddg-pharmfac.net/AllerTOP/index.html) was used [54]. Lastly, the toxicity screening was performed using ToxinPred (https://webs.iiitd.edu.in/raghava/toxinpred/algo.php) to finalise B-cell epitopes [55].

### 5.3. Prediction of Cytotoxic T-Lymphocyte epitope with MHC-I binding

For the prediction of Cytotoxic T-Lymphocyte (CTL) or CD8^+^ T-cell epitope, an online epitope prediction tool NetCTL v1.2 server was used [56]. The prediction model for epitope binding affinity, proteasomal C-terminal cleavage, and TAP (transporter-associated antigen processing) was established with specific cutoff values. The cutoff values used were 0.05 for binding affinity prediction, 0.15 for proteasomal C-terminal cleavage, and 0.75 for TAP transport efficiency. The predicted epitopes were then evaluated to check the MHC-I binding ability using the IEDB MHC-I Binding Predictions server (http://tools.immuneepitope.org/mhci/) [57].

### 5.4 Prediction of Helper T-Lymphocyte epitope with MHC-II binding

For the prediction of Helper T-Lymphocyte (HTL) or CD4^+^ T-cell epitope, IEDB MHC class-II Binding Predictions server (http://tools.immuneepitope.org/mhcii/) was used [58]. For each antigenic protein, epitopes were predicted depending on the IC_50_ score, which indicates the affinity of peptides towards the MHC-II alleles. The IC_50_ value less than 50 nM corresponded to the highest binding affinity towards MHC-II, the IC_50_ value less than 500 nM suggested midrange affinity, while the IC_50_ value less than 5000 nM indicated the lowest binding affinity.

### 5.5 Evaluation of antigenicity, allergenicity, and toxicity of predicted CTL and HTL

Antigenicity of all the predicted CTL and HTL were checked using VaxiJen v2.0, taking 0.4 as the threshold value. Subsequently, the IFNepitope server (https://crdd.osdd.net/raghava/ifnepitope) was utilized to screen the CD8^+^ and CD4^+^ epitopes to evaluate the capability to induce interferon-γ (IFN-γ) [59]. Furthermore, the allergenicity of the CD8^+^ and CD4^+^ epitopes were screened with the AllerTop v2.0 server & ToxinPred server was used for the prediction of toxicity for the final selection of the epitopes.

### 5.6 Construction, modelling and validation of Multi-Epitope Vaccine (MEV)

The predicted CD8^+^ and CD4^+^ T-cell epitopes, which were chosen for the vaccine construction based on factors such as high antigenicity, non-allergenicity, and non-toxicity, were joined to create the final multi-epitope vaccine. Three experimentally obtained B-cell epitopes (TRMQTQINGLNQGVSNAND, INSAADDAAGLAIATR, QLAVQASNGPLSASDASALQQEVAQQISEVNR) from **BPSL3319** were also used in MEV construction [60]. Because OppA from *B. pseudomallei* has demonstrated effectiveness as a vaccine candidate, it was chosen as the scaffold. Flexible non-immunogenic portions of OppA were removed by truncating OppA, and the chosen epitopes were carefully positioned to design the chimeric protein. In order to create a compact, spherical tertiary structure, an iterative process of construction, deconstruction, and model improvement was used. AAY linkers were used to join CTL epitopes, whereas GPGPG linkers were used to join HTL epitopes. Using the EAAAK linker, human β-defensin-3 adjuvant (the 45-amino acid peptide, recognized for its immunomodulatory qualities) was fused to the N-terminal of the CTL vaccine to improve the immunological response. Afterwards, AllerTop v2.0 and VaxiJen v2.0 were used to evaluate the vaccine design for allergenicity and antigenicity, respectively. An AI-based algorithm, Robetta, was used to predict the 3D structure of the designed MEV. Then, the Galaxy Refinement server was used to improve the structure quality. Furthermore, the energy-minimized, predicted structure was evaluated using PROCHECK [61], and the Ramachandran plot was obtained. CABS-Flex 2.0 server [62] was employed for the assessment of the flexibility of the designed vaccine construct.

### 5.7 Analysis of MEV-TLR2 Interaction using Molecular Docking

It was found that TLR2 significantly modulates the host immune response in *B pseudomallei* infection [22]. For this reason, to further study the interaction, it was decided to perform molecular docking studies between the MEV construct and TLR2. For the molecular docking studies, the ClusPro v2.0 server was used [63]. Computational steps like rigid body docking, clustering based on Root-Mean-Square Deviation (RMSD), and structural refinement using energy minimization were used to predict the docked conformation. The lowest energy score and binding efficacy were considered to select the best-docked complex. The docked MEV-TLR2 complex was subjected to Binding Free Energy calculation by the MM/GBSA method using the Hawkdock server (http://cadd.zju.edu.cn/hawkdock/) [64].

### 5.8 Analysis of conformational dynamics of MEV and the MEV-TLR2 complex

For the assessment of the conformational dynamics of the MEV, molecular dynamics (MD) simulation studies were performed using the GROMACS 5.1.2 platform (https://www.gromacs.org) [65]. Protein topology was generated using the Charmm27 force field [66]. In order to solvate the simulation system, TIP3P water box was used, followed by energy minimization to avoid steric clashes. To address the long-range electrostatic interactions, the particle mesh Ewald (PME) approach with a 12Å cut-off was used to equilibrate the system utilizing the NVT and NPT ensemble. A 50 ns simulation was performed, and trajectories were examined using the built-in scripts. After the MD simulation was successfully completed, water and ions were eliminated from the trajectory, and the periodicity was rectified. The adjusted trajectory was used to compute the root mean square deviation (RMSD), root mean square fluctuation (RMSF), and the radius of gyration (Rg). A similar approach was used to assess the trajectories obtained from MD simulation of the MEV-TLR2 complex.

### 5.9 Immune simulation of constructed MEV

For the evaluation of antibody response after the administration of the MEV construct, the C-IMMSIM server (http://kraken.iac.rm.cnr.it/C-IMMSIM/) [67] was used. Information on the activation of humoral and cellular defences, including B-cells, dendritic cells, Natural Killer (NK) cells, Helper T lymphocytes (HTL), Cytotoxic T lymphocytes (CTL), immunoglobulins and cytokines, following vaccination delivery, was provided by the C-IMMSIM server. For most commercial vaccinations, there is a minimum suggested interval of 4 weeks between doses 1 and 2. The immunological responses were assessed using 3 injections administered at 4-week intervals, which were translated into 1, 84, and 168 time steps (1 time step is equivalent to 8 hours in real life). Each dosage comprised 1000 vaccine particles, and the simulation was run for 1050 time steps (350 days).

### 5.10 Codon optimization & *In-silico* cloning of the MEV

The DNA sequence of the constructed MEV was obtained using the Reverse Translate Tool. In order to express the multi-epitope vaccine in the E. coli (K12 strain) host, codon optimization of the final vaccine construct was performed utilizing the JCat tool [68]. Furthermore, the following options: ‘avoid the rho-independent transcription termination’, ‘avoid prokaryote ribosome binding site’, and ‘avoid restriction enzymes cleavage sites’, were chosen for the codon optimization. The JCat output takes account of the codon adaptation index (CAI) and GC content (%), these parameters provide information for the assessment of protein expression levels. The CAI score of 1.0 is considered ideal, though a score higher than 0.8 is also considered as a good score. The ranges of GC content should lie between 30% and 70%, and the values outside that range indicate adverse effects on the efficiencies of transcription & translation. For *in-silico* cloning of the optimized gene sequence of the final vaccine construct, the pET-28a (+) vector was chosen having the *BamHI* and *Xho*I restriction sites added. To ensure the successful expression of the vaccine, SnapGene tool (https://www.snapgene.com) was used to insert the optimized DNA sequence into the pET-28a (+) vector, including the restriction sites. The SOLpro tool was utilized to predict the solubility of MEV [69].

## AUTHOR CONTRIBUTIONS

DH and AR conceptualized the project, designed the experiments, analysed the data, designed the MEVs, prepared the figures and edited the manuscript. SR performed the experiments data analysis MD simulation and wrote the manuscript. AD prepared figures, preliminary analysis and wrote the manuscript. AKD provided the expertise and infrastructure required to run MD simulations.

## ACKNOWLEDGEMENT

SR acknowledges IIT Kharagpur for institute fellowship. AD acknowledges IIEST Shibpur for institute fellowship. AKD would like to acknowledge the Department of Biotechnology, Government of India (GOI), File no. BT/PR40990/MED/29/1540/2020 for funding. DH acknowledges St. Xavier’s College (Autonomous), Kolkata for Intramural Research Grant (IMSXC2023-24/007). AR acknowledges research grant from WB-DST (2149/STBT/13015/5/2023/WBSCST SEC).

