## Supplementary for "Designing a novel Scaffold-Based Multi-Epitope Vaccine to Combat Melioidosis Caused by Burkholderia pseudomallei: An *In-silico* and Immunoinformatics approach"

### Figure Legend:

**Figure 1:** Representation of the MEV secondary structure. The β-strand regions are depicted by yellow colour, the α-helix regions are depicted by pink colour and the grey colour indicates the coil regions.

**Figure 2:** Representation of Ramachandran Plot indicates that 90.2% of the amino acids are in the most favoured regions, 7.5% in the additionally allowed,1.3% in the generously allowed, and 0.9% in disallowed regions.

**Figure 3:** Codon Optimized DNA sequence obtained using the JCat Tool.

**Figure 1**

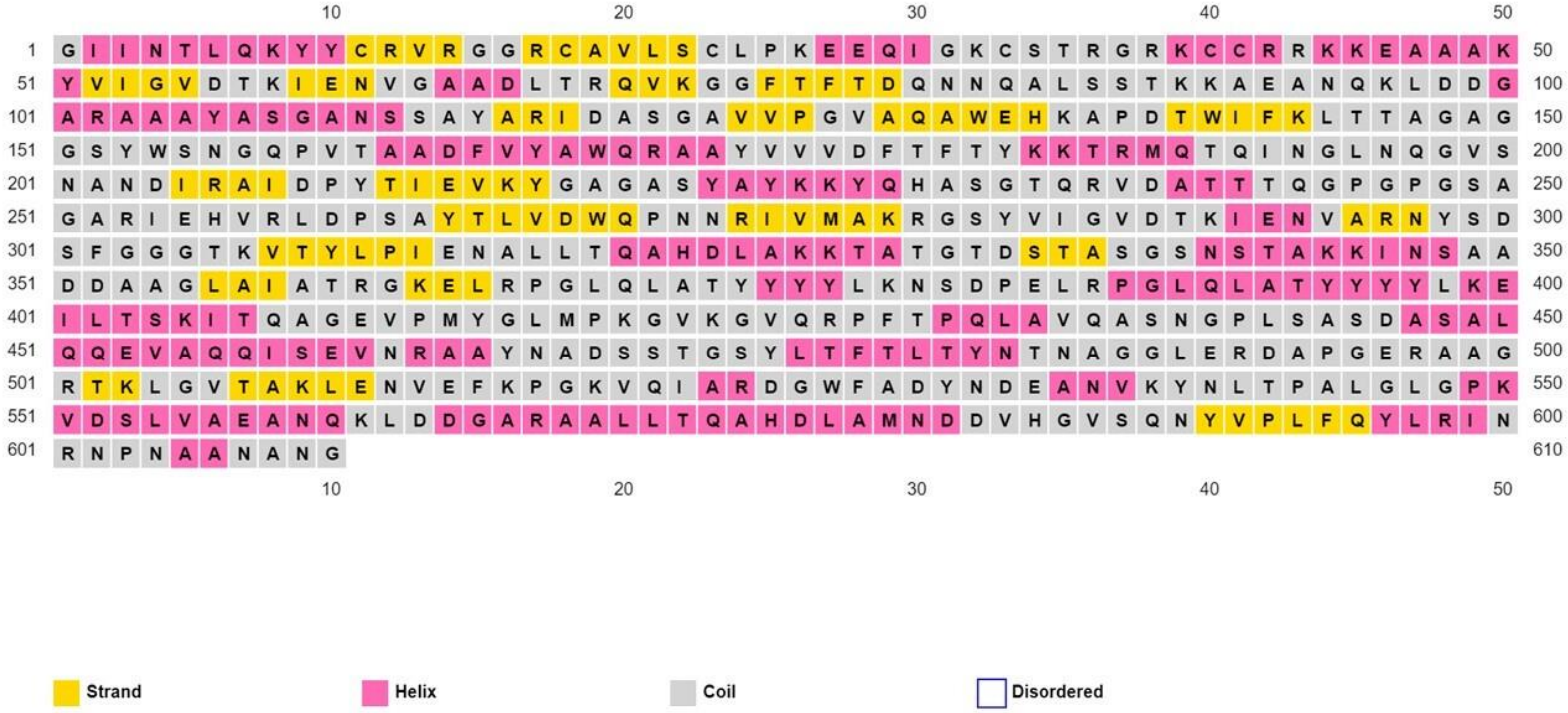
10 20 30 40 50
1 G I I N T L Q K Y Y C R V R G G R C A V L S C L P K E E Q I G K C S T R G R K C C R R K K E A A A K 50
51 Y V I G V D T K I E N V G A A D L T R Q V K G G F T F T D Q N N Q A L S S T K K A E A N Q K L D D G 100
101 A R A A A Y A S G A N S S A Y A R I D A S G A V V P G V A Q A W E H K A P D T W I F K L T T A G A G 150
151 G S Y W S N G Q P V T A A D F V Y A W Q R A A Y V V V D F T F T Y K K T R M Q T Q I N G L N Q G V S 200
201 N A N D I R A I D P Y T I E V K Y G A G A S Y A Y K K Y Q H A S G T Q R V D A T T T Q G P G P G S A 250
251 G A R I E H V R L D P S A Y T L V D W Q P N N R I V M A K R G S Y V I G V D T K I E N V A R N Y S D 300
301 S F G G G T K V T Y L P I E N A L L T Q A H D L A K K T A T G T D S T A S G S N S T A K K I N S A A 350
351 D D A A G L A I A T R G K E L R P G L Q L A T Y Y Y Y L K N S D P E L R P G L Q L A T Y Y Y Y L K E 400
401 I L T S K I T Q A G E V P M Y G L M P K G V K G V Q R P F T P Q L A V Q A S N G P L S A S D A S A L 450
451 Q Q E V A Q Q I S E V N R A A Y N A D S S T G S Y L T F T L T Y N T N A G G L E R D A P G E R A A G 500
501 R T K L G V T A K L E N V E F K P G K V Q I A R D G W F A D Y N D E A N V K Y N L T P A L G L G P K 550
551 V D S L V A E A N Q K L D D G A R A A L L T Q A H D L A M N D D V H G V S Q N Y V P L F Q Y L R I N 600
601 R N P N A A N A N G 610
10 20 30 40 50
Strand
Helix
Coil
Disordered

**Figure 2**

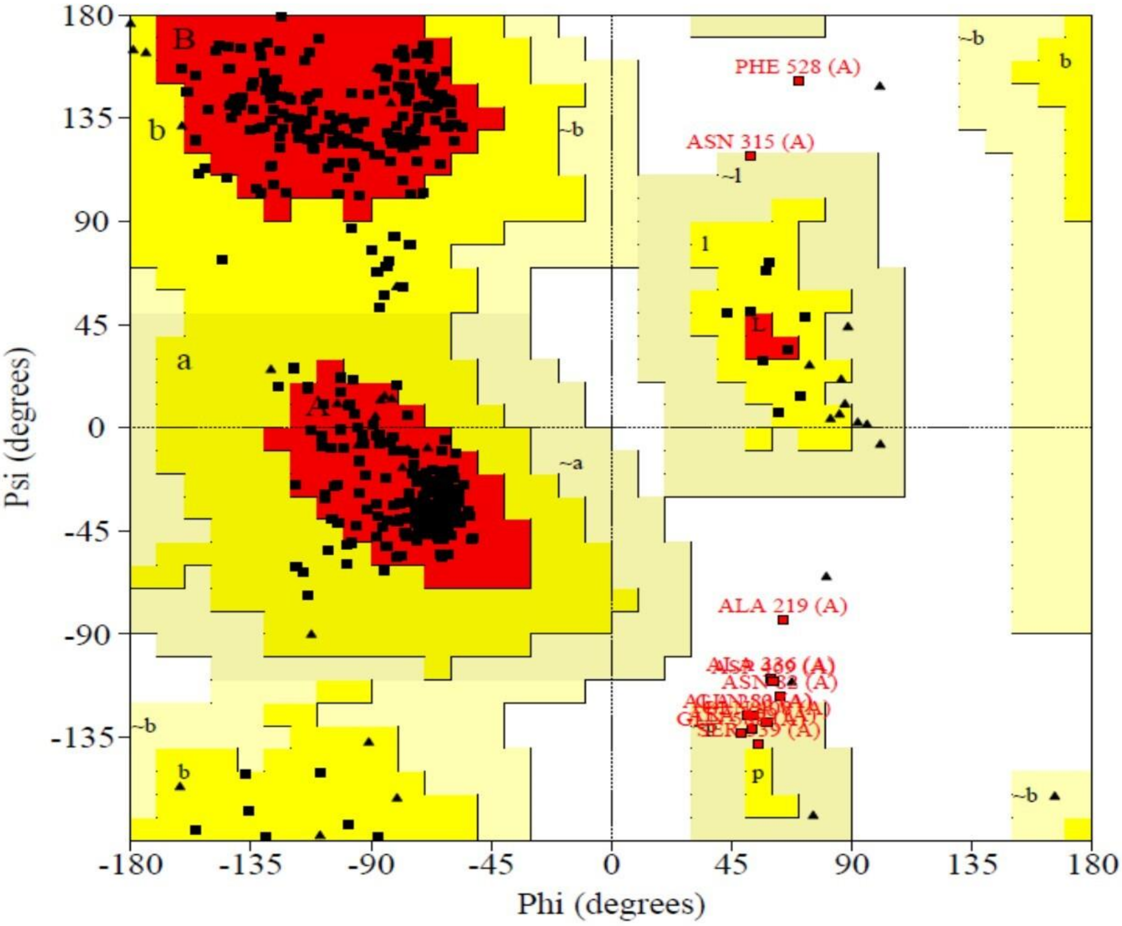

PHE 528 (A)
ASN 315 (A)
ALA 219 (A)
Psi (degrees)
Phi (degrees)
180
135
90
45
0
-45
-90
-135
-180
-135
-90
-45
0
45
90
135
180


**Figure 3**

**Codon Optimized DNA sequence**

>filtered DNA sequence consisting of 1830 bases

GGTATCATCAACACCCTGCAGAAATACTACTGCCGTGTTCGTGGTGGTCGTTGCGCTGTT
CTGTCTTGCCTGCCGAAAGAAGAACAGATCGGTAAATGCTCTACCCGTGGTCGTAAATGC
TGCCGTCGTAAAAAAGAAGCTGCTGCTAAATACGTTATCGGTGTTGACACCAAAATCGAA
AACGTTGGTGCTGCTGACCTGACCCGTCAGGTTAAAGGTGGTTTCACCTTCACCGACCAG
AACAACCAGGCTCTGTCTTCTACCAAAAAAGCTGAAGCTAACCAGAAACTGGACGACGGT
GCTCGTGCTGCTGCTTACGCTTCTGGTGCTAACTCTTCTGCTTACGCTCGTATCGACGCT
TCTGGTGCTGTTGTTCCGGGTGTTGCTCAGGCTTGGGAACACAAAGCTCCGGACACCTGG
ATCTTCAAACTGACCACCGCTGGTGCTGGTGGTTCTTACTGGTCTAACGGTCAGCCGGTT
ACCGCTGCGGACTTCGTGTACGCGTGGCAGCGTGCTGCTTACGTTGTTGTTGACTTCACC
TTCACCTACAAAAAAACCCGTATGCAGACCCAGATCAACGGTCTGAACCAGGGTGTTTCT
AACGCTAACGACATCCGTGCTATCGACCCGTACACCATCGAAGTTAAATACGGTGCTGGT
GCTTCTTACGCTTACAAAAAATACCAGCACGCTTCTGGTACCCAGCGTGTTGACGCTACC
ACCACCCAGGGTCCGGGTCCGGGTTCTGCTGGTGCTCGTATCGAACACGTTCGTCTGGAC
CCGTCTGCTTACACCCTGGTTGACTGGCAGCCGAACAACCGTATCGTTATGGCTAAACGT
GGTTCTTACGTTATCGGTGTTGACACCAAAATCGAAAACGTTGCTCGTAACTACTCTGAC
TCTTTCGGTGGTGGTACCAAAGTTACCTACCTGCCGATCGAAAACGCTCTGCTGACCCAG
GCTCACGACCTGGCTAAAAAAACCGCTACCGGTACCGACTCTACCGCTTCTGGTTCTAAC
TCTACCGCTAAAAAAATCAACTCTGCTGCTGACGACGCTGCTGGTCTGGCTATCGCTACC
CGTGGTAAAGAACTGCGTCCGGGTCTGCAGCTGGCTACCTACTACTACTACCTGAAAAAC
TCTGACCCGGAACTGCGTCCGGGTCTGCAGCTGGCTACCTACTACTACTACCTGAAAGAA
ATCCTGACCTCTAAAATCACCCAGGCTGGTGAAGTTCCGATGTACGGTCTGATGCCGAAA
GGTGTTAAAGGTGTTCAGCGTCCGTTCACCCCGCAGCTGGCTGTTCAGGCTTCTAACGGT
CCGCTGTCTGCTTCTGACGCTTCTGCTCTGCAGCAGGAAGTTGCTCAGCAGATCTCTGAA
GTTAACCGTGCTGCTTACAACGCTGACTCTTCTACCGGTTCTTACCTGACCTTCACCCTG
ACCTACAACACCAACGCTGGTGGTCTGGAACGTGACGCTCCGGGTGAACGTGCTGCTGGT
CGTACCAAACTGGGTGTTACCGCTAAACTGGAAAACGTTGAATTCAAACCGGGTAAAGTT
CAGATCGCTCGTGACGGTTGGTTCGCTGACTACAACGACGAAGCTAACGTTAAATACAAC
CTGACCCCGGCTCTGGGTCTGGGTCCGAAAGTTGACTCTCTGGTTGCTGAAGCTAACCAG
AAACTGGACGACGGTGCTCGTGCTGCTCTGCTGACCCAGGCTCACGACCTGGCTATGAAC
GACGATGTTCACGGTGTAAGCCAGAACTACGTTCCGCTGTTCCAGTACCTGCGTATCAAC
CGTAACCCGAACGCTGCTAACGCTAACGGT
